# No evidence for the positive relationship between genetic correlations and heritabilities

**DOI:** 10.1101/039388

**Authors:** Szymon M. Drobniak, Mariusz Cichoń

**Author notes:** Corresponding author, Institute of Enviornmental Sciences, ul. Gronostajowa 7, 30-387 Kraków, Poland.

## Abstract

Quantitative genetics predicts, that traits subjected to strong selection should show low heritability and may yield biased estimates of genetic correlations (rg). Similar pattern may also appear if genetic sources of variation are confounded with non-genetic sources. Thus, a positive relationship between genetic correlations and heritabilities (h^2^) of underlying traits might be observed. Here we test this prediction using a large dataset of published estimates of genetic correlations and employing a powerful meta-analytical approach. We considered both between-traits and cross-sex genetic correlations. We failed to find support for the prediction about a positive r_g_ – h^2^ relationship: our analysis based on nearly 1000 published estimates of genetic parameters indicates that the predicted relationship is weak and statistically non-significant. Thus, low heritability does not preclude the possibility of detecting substantial genetic correlations. Our meta-analysis indicates that published estimates of genetic parameters coming from various experimental designs and obtained using different statistical techniques are not significantly biased in case of weakly-heritable traits.

## Introduction

Estimation of genetic parameters such as heritability or evolvability became a standard procedure in most ecological and evolutionary studies (Steppan et al. 2002; L. E. B. Kruuk and JD Hadfield 2007; Hansen et al. 2011). With the advent of modern statistical and computational tools, large, sparse and unbalanced datasets typical for studies performed in wild populations became readily available in quantitative genetics. However, phenotypic traits are not separate entities and usually it is desired to analyze them together with other traits, seeking for multivariate genetic patterns (e.g. Steppan et al. 2002; Jensen et al. 2003; Colautti et al. 2010; King et al. 2011). In this sense, genetic correlations lie in the very core of quantitative evolutionary genetics. Recently studies reporting genetic correlations in natural populations became numerous. However, field estimates of genetic parameters obtained from unplanned breeding events, using limited sample sizes available in the field might be not very accurate (Kruuk 2004).

Genetic correlation describes the degree to which two traits share their genetic background, i.e. the extent to which the two traits are influenced by the same set of genes (Lynch and Walsh 1998). Mathematically it can be defined as 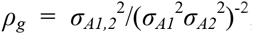, where indexes A1 and A2 indicate respective additive genetic variances, whereas A1,2 describes additive genetic covariance. Importantly, genetic correlation may be defined not only between different traits but also for the same trait between different classes of individuals (Lynch and Walsh 1998). In that sense one may consider cross-sex *rg*, genotype-by-environment *r_g_* or longitudinal *r_g_* (genotype-by-age interaction). Within-trait measures of genetic association are of key interest as they serve as measures of various genotypic interactions. Patterns of within-trait genetic correlations appear to be more complex than expected earlier and may be present as general, intrinsic properties of evolving biological systems (see Hoffmann and Merilä 1999; Brommer et al. 2007; Poissant et al. 2010 for reviews and examples of within-trait *r_g_* studies).

The true evolutionary significance of genetic correlations is summed up in the breeders’ equation (Roff 2006):

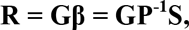

where **R** denotes change in the trait value due to selection, **G** is the genetic variance-covariance matrix, **β** is the vector of direct (univariate) selection gradients, **P** is the phenotypic (co)variance matrix and **S** stands for selection differentials. Calculating evolutionary change in, say, the first trait yields 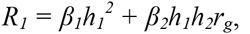 where *h_i_* stands for the square-root of the respective heritability (denoted as 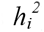) (Roff 2006). Constraint imposed by this equation means, that in the presence of strong genetic correlation between the two traits, any selective change in the second trait will be reflected in the first trait leading to correlated evolutionary change (Lynch and Walsh 1998; Roff 2006). On the other hand, opposing fitness optima for the two traits will lead to zero or little net evolutionary change, a process called inter-locus genetic conflict (Andres and Morrow 2003). Studies exploring patterns of correlated evolution of quantitative traits are numerous. Less understood, however, are mechanisms that actually maintain bonds of genetic correlations. We still lack answers to problems such as the origin of genetic correlations or their relationship to other genetic parameters. The latter should be of particular interest as it would not only provide interesting insights into evolutionary genetic processes but also offer and efficient tool for validating reliability and robustness of quantitative genetic studies.

The prediction that genetic correlations should be related (either in the sense of the magnitude of the effect or it’s detectability) to heritability/additive genetic variance has no mathematical ground. Although correlation is directly derived using respective variances (i.e. the components of heritability), covariance is the main source of correlation and, mathematically, covariance does not depend on the variance (and hence on heritability; Lynch and Walsh 1998; Quinn and Keough 2002). However, several statistical and genetic phenomena might be regarded as possible generators of such relationship. Firstly, as proposed by (Ronald A. Fisher 1930) and Price and Schluter (1991), traits more closely related to fitness (such as direct proxies of fitness and life-history traits) should have lower heritability than e.g. morphological or physiological traits. In fact, such patterns has been reported in a number of studies (e.g. Gustafsson 1986; Merilä and Sheldon 1999; Kruuk et al. 2000; Teplitsky et al. 2009). Fisher (1930) argued that low heritabilities of such traits should result from lack of genetic variability in fitness in populations at evolutionary equilibrium. These findings were further developed by Price and Schluter (1991) who concluded, that low heritability of life-history traits might result from environmental variability added to their variance by other metric (e.g. morphological) traits closely related to life-history traits. Importantly, not only genetic variance but also genetic covariance between life-history traits could be biased due to environmental sources of variability, yielding low levels of genetic correlations between life-history characters. Therefore it is possible, that such patterns present in life-history traits could be responsible for putative *r_g_* – *h^2^* correlations.

Another possible source of spurious correlations between *h^2^* and *r_g_* could arise due to statistical sampling issues. Estimation of genetic correlations is demanding both in terms of sample sizes and statistical methodology (Lynch and Walsh 1998). Although sampling distribution of genetic correlations is approximately normal (Fisher 1928), lower sample sizes result in much higher variance levels and wider confidence intervals (Brown 1969). Also, for some statistical procedures such as parent-offspring regression, lower sample sizes may be associated with biased estimates of standard errors of *r_g_* (Lynch and Walsh 1998), making such genetic correlations more difficult to detect. As standard errors for heritabilities should suffer similarly from low sample sizes used in their estimation (Lynch and Walsh 1998), this might be the source for the presumed *r_g_* – *h^2^* correlation. The problem complicates if one considers additional, non-genetic sources of variation in phenotypic traits. If statistical procedures fail to separate strong common environmental or parental effects from genetic sources of variation, one may end up estimating genetic correlations with genetic and non-genetic sources of variation confounded. Patterns of genetic covariation may become blurred in such situations, leading to biased estimates of genetic correlations. Lower heritabilities would suffer more as they would be obscured by additional variance to a greater extent – hence generating assumed relationship between *h^2^* and r_g_.

Finally, expected values of *r_g_* may depend on the heritabilities of traits under strong directional selection (Lande and Price 1989). As simulation studies indicate, strong selection operating on one trait may severely bias *r_g_* and the degree of bias depends on the heritability of the trait under selection. In particular it was proved that covariance between trait 1 in parents and trait 2 in offspring *Cov_s_*, after selection acting on parents with respect to the trait 1, equals:

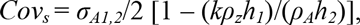

where *σ_A1,2_* – additive genetic covariance, *k* – strength of selection, *ρ_z_* – phenotypic correlation before selection, *h_i_* – square-root of heritability, *ρ_A_* – genetic correlation (Lande and Price 1989). It can be seen from this equation that selection may generate spurious relationships between estimated covariance (and hence correlation) and heritability.

The abovementioned considerations are probably partially responsible for a widespread opinion that weakly heritable traits should exhibit lower levels of genetic correlations. Surprisingly, we did not find any published evidence for such relationship. Thus, the validity and prevalence of such predictions remains to be discovered. In this study we explicitly explore the relationship between genetic correlations and heritabilities, both theoretically and using meta-analytical approach on empirical published data. We test the prediction that genetic correlations are positively associated with heritabilities of underlying traits.

## Materials and methods

### Data collection and preparation

We searched publicly available databases using the following keywords: “genetic correlation”, “genetic covariance”, “heritability”. All papers returned by our queries were classified into two groups: the first set contained papers analyzing genetic correlations between different traits (dataset X), the second one considered only cross-sex genetic correlations (dataset S). We decided to subdivide our analysis into two sets of data taken different features of cross-traits and cross-sex genetic correlations, both from the point of view of their biological relevance and statistical properties (i.e. different null hypotheses tested in each of these two categories; see Discussion). All papers were scanned for the estimates of genetic correlations and heritabilities. Papers were retained for further analyses only if they provided estimates of both heritabilities for traits underlying *r_g_* and standard errors or confidence intervals for the estimates of genetic correlations. Studies of cross-sex genetic correlations were verified with the recent meta-analysis on sexual dimorphism (Poissant et al. 2009) and hence our study contains a similar set of papers, updated with respect to papers published after 2009. As for the cross-trait genetic correlations we decided include only studies dated back to 2009. Our motivation for that was a huge number of published estimates of genetic correlations, most of them from agricultural studies, which quickly outnumbered the number of estimates in the first dataset (S). In our opinion it did not bias our estimates since with the increasing number of studies the mean effect size asymptotically approaches a single value (Fig. 1).

Our dataset contained the following variables: genetic correlation, heritability of the first and second trait, standard error for genetic correlation, significance of the *r_g_* at the 95% level (binary variable: significantlnon-significant; variable obtained using provided standard errorslconfidence intervals), trait type of the first and second trait (categorical variable: morphology, physiology, behavior, developmental, life-history, fitness), type of the study (agricultural, laboratory, field), type of the statistical procedure applied (full-sib design, half-sib design, animal model, parent-offspring regression, ANOVA). To account for the lack of independence of different genetic correlations we included additional categorical variables: species (and if possible breed), study id (equal to the bibliographical record for the study), trait 1 and 2 id (relevant for cross-trait genetic correlations where different traits might appear several times in different combinations).

**Fig. 1.**
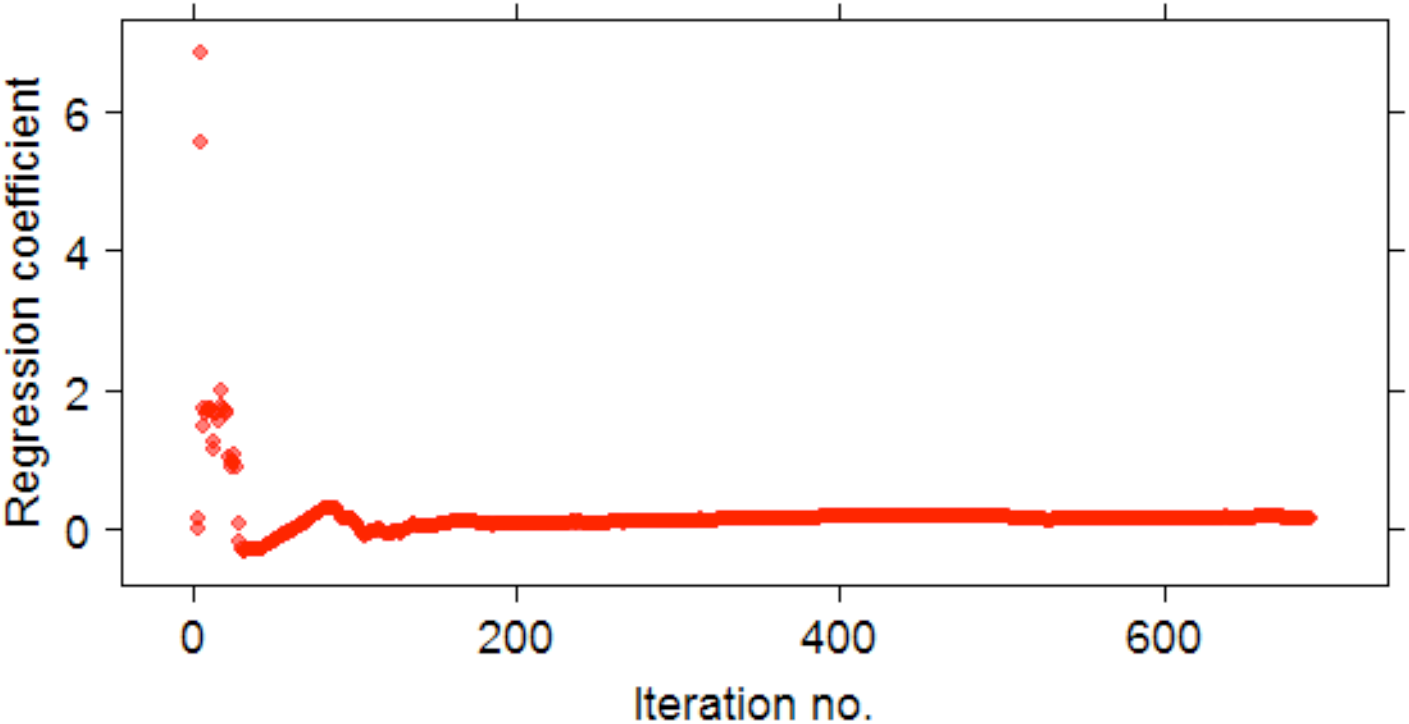
Estimates of the regression coefficient for the *r_g_* – *h^2^* relationship after including subsequent rows of data, beginning with the newest studies. Iteration number represents the number of included studies.

Wherever possible, original standard errors where used. However, if only confidence intervals were provided, we calculated approximate standard errors of the estimates dividing half of the confidence interval by the value of the appropriate quantile of the standard normal distribution (which equals 1.96 for the 95% confidence interval).

Effect size in our meta-analysis was expressed as the square-root of the modulus of Fisher’s Z-transformed correlation coefficient (√|Z|), where *Z*=ln[(1+*r_g_*)l(1-*r_g_*)]. The choice of Fisher’s Z is natural when analyzing correlations (Borenstein et al. 2009). However, as we were mainly interested in the overall magnitude of genetic correlation and not in its sign, we decided to use the modulus (or absolute value) of *Z*. *Z* is approximately normally distributed and thus our |*Z*| variable would have a half-normal asymmetrical distribution. To overcome this we applied a normalizing transformation and hence our effect size took the form of √|*Z*|.

Heritabilities of specific traits were assigned arbitrarily to the first and the second independent variable as they appear in the analysed paper. Thus, to avoid any possible bias related to arbitrary allocation of the heritabilities to respective independent variables and to reduce complexity of the model we decided to use average heritability (arithmetic mean) as the independent variable. Additionally, the absolute difference between respective heritabilities was included in all models to account for possible differences in *h^2^* between traits. To ensure that averaging heritabilities did not introduce any bias into our estimates we generated 1000 estimates of regression coefficients for the *r_g_* – *h^2^* relationship using both heritabilities instead of the average *h^2^*, each time randomly changing the assignment of both heritabilities to the first and the second independent variable. Distributions of such regression coefficients largely overlapped the distributions of coefficients expected under the null hypothesis of no correlation, hence confirming that arbitrary order of heritabilities wouldn’t introduce any bias (Fig. 2).

**Fig. 2.**
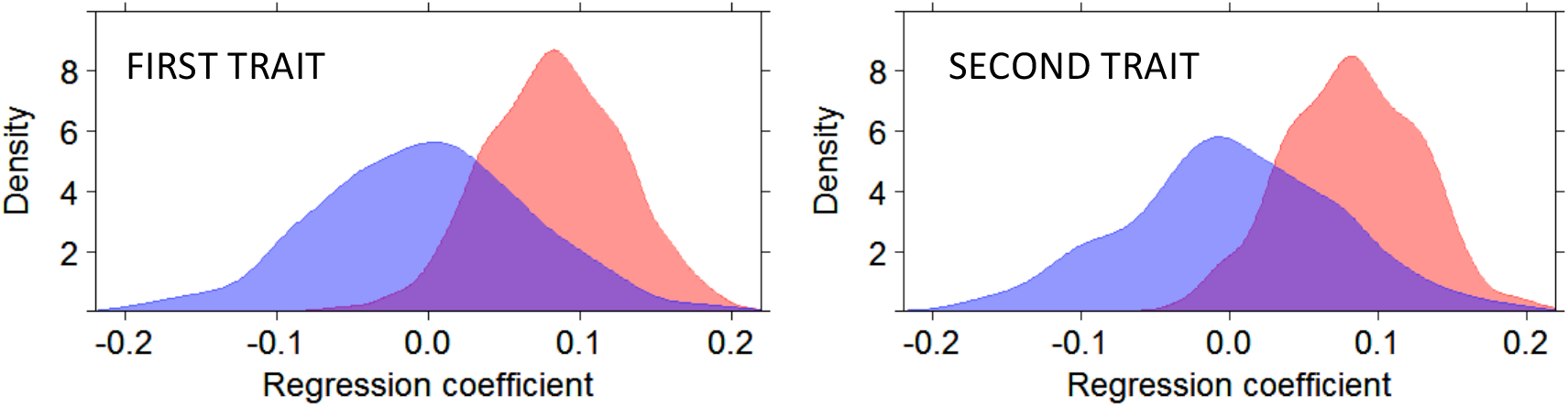
In red: distribution of the regression coefficients for the relationship between effect size and average *h^2^* for models with randomized order of traits (i.e. heritabilities of trait 1 and trait 2 were swapped, N=1000). In blue: distributions of the estimates of regression coeffcients obtained by ordinary randomization test, without re-ordering trait’s heritabilities.

In total 60 studies reporting 691 values of *r_g_* were included in the dataset X and 50 studies reporting 219 estimates of *r_g_* were included in the dataset S. The full set of analyzed data plus references to all analyzed studies is provided in the electronic supplementary materials.

### Statistical analyses

As explained above, cross-sex (dataset S) and cross-trait (dataset X) genetic correlations were analyzed separately and thus two sets of models are considered below, differing with the respect of the type of correlation. The square-root of |*Z*| was treated as the response in both models and the following independent fixed variables were defined: average *h^2^*, difference between respective *h^2^*’s, trait type, statistical methodology and study type (see explanations above). Additionally, species, study ID and traits ID’s (the latter only for the cross-trait analyses) were included as random effects. We considered all possible interactions of the categorical variables with both covariates (average *h^2^* and of *h*^2^’s difference). However, all interactions and all categorical variables appeared non-significant (P>0.3 in all cases) indicating that both the magnitude of *r_g_* and the slope of the *r_g_* – *h^2^* relationship were uniform across different trait types, statistical procedures and study types. Hence, we decided to remove these interactions and categorical variables from our models.

Additionally, a generalized linear mixed model with binomial error structure and logit link function was used to analyze whether *h^2^* affects the probability of obtaining significant estimates of *r_g_*. This model contained similar fixed and random effects. Because for cross-sex genetic correlations an appropriate null hypothesis (H_0_: *ρ_g_* = 1) is equal to the boundary of the space of possible values, standard errors should not be used to test this hypothesis (the use of likelihood ratio tests or information theory is preferred here; see Fox and Wolf 2006). Thus, we did not analyze significance of cross-sex *r_g_*’s in that way but only significance of cross-trait genetic correlations (where the appropriate null hypothesis is of the form H_0_: *ρ_g_* = 0).

All models were run as meta-analytical linear models and thus they accounted for the sampling error of published estimates of genetic correlations. The typical linear mixed model has the following form: **y** = **Xβ** + **Zu** + **e**, where **X** and **Z** are incidence matrices for fixed and random effects, **β** is the vector of fixed effects coefficients, **u** ∼ *N*(0, **G**) is the vector of random effects, **e** ∼ *N*(0, **R**) is the vector of error terms. In a random-effects meta-analysis another term comes in: **y** = **Xβ** + **Zu** + **m** + **e**, where **m** is the vector of error measurements (e.g. standard errors as in our case).

In order to determine possible sources of bias in meta-analyses, we employed funnel graphs by plotting residual effect size (residuals from models fitted to Fisher’s *Z* effect size) against the reciprocals of standard errors (precision of estimates). Egger’s regression was applied to funnel plots (Egger et al. 1997). This method estimates possible asymmetry of funnel plots by testing the significance of the linear slope fitted to the funnel plot; a slope significantly different from zero indicates possible publication bias. We used Egger’s regression on residual effect sizes rather than raw effect sizes in order to account for non-independence of data points due to complex random effects structure (Horváthová et al. 2012).

Analyses were performed in R (2.14.2; R Development Core Team 2011). Meta-analytical linear and generalized linear mixed models were fit using MCMCglmm package (Hadfield 2010). All models were run for 2**x**10^6^ iterations, first 50000 iterations were discarded and samples were drawn from posterior every 100^th^ iteration. Priors for all random effects were set as weak half-Cauchy distributions (with parameters V=1, nu=0.002, alpha.mu=0, alpha.V=1000). Priors for residual variance were set as inverse-Wishart distributed (with parameters V=1, nu=0.002). All models were checked for autocorrelation issues by visual inspection of time-series plots, however no problems were detected.

### Analysis of simulated correlations

To search for possible sources of the *r_g_* – *h^2^* relationship we sampled toy data from bivariate normal distributions (*BVN*), diversifying parameters used in each sampling. Our aim was to determine the influence of both additive genetic variance *V_A_* (and hence heritability) and the presence of additional non-genetic variance (confounded with V_A_) on the estimates of r_g_. All data were sampled from *BVN* using the rmvnorm procedure from mvtnorm library. All samplings were made using distributions with zero mean **μ** = [0, 0] with the population value of genetic correlation being *ρ_G_* = 0.8. We used 13 decreasing values of additive genetic variance (ranging from 0.05 to 2.0) and fro each of these values set the additive genetic (co)variance matrix in order to maintain constant genetic correlation of 0.8. We performed six series of samplings for all values of *V_A_*’s, each time increasing the value of additional, non-genetic values with variance 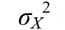 (sampled from a univariate normal distribution with mean zero and variance 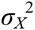 and thus introducing no additional correlation; 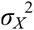 ranged from 0.05 to 0.5). Thus, 78 samplings were performed for each combination of *V_A_* and 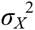. For each combination, the estimated value of *r_g_* was obtained using the cor.test function. Estimated genetic correlations for all six series of samplings (according to values of 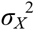) were then plotted against *V_A_* values used to sample from BVN. We assumed that the additional variance 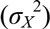 is not distinguishable from the true population value of *V_A_* and hence values used generating the abovementioned plot were 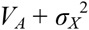.

## Results

The collected data provide a wide range of *r_g_* and *h* values. In general, cross-sex genetic correlations were mostly positive and close to unity, which could be expected taken their biological properties. On the other hand, between-traits genetic correlations were represented mainly by low, close-to-zero values (Fig. 3). Interestingly, the distributions of cross-sex and cross-trait heritabilities were different: the former was symmetrical with the mode around 0.5, the latter was clearly bimodal, indicating that high and moderate to low heritabilities were the most common (Fig. 3).

**Fig. 3.**
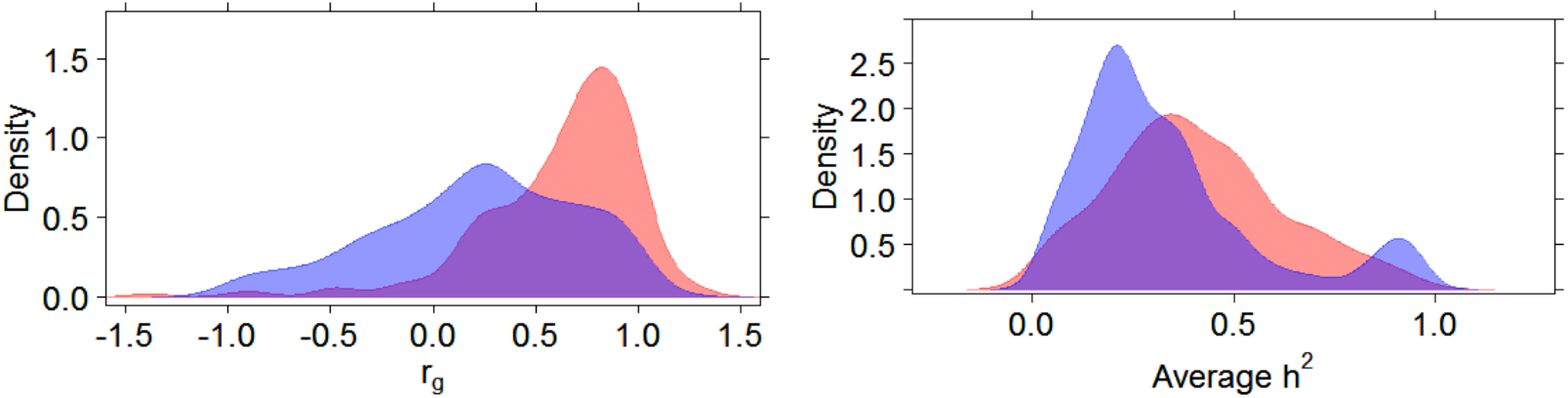
Left: distributions of cross-trait (blue) and cross-sex (red) genetic correlations. Right: distributions of of heritabilities for cross-sex (red) and cross-trait (blue) datasets.

Analysis of the absolute Fisher’s Z values (√|*Z*|) indicated that the magnitude of genetic correlation is not significantly correlated with the average heritability of underlying traits (Fig. 4 and 5). A relationship between √|*Z*| and mean heritability was stronger in case of cross-sex genetic correlations compared to cross-trait *r_g_*’s, however it remained marginally non-significant (Table 1). The size of the effect (i.e. the regression coefficient for the relationship between genetic correlation and average *h^2^*) was low for both datasets (Table 1, Fig. 4 and 5). In case of cross-sex genetic correlations taxon explained significant proportion of variance in effect sizes (Table 1), whereas in between-trait genetic correlations, most variance was explained by the trait ID. Nevertheless, posterior credibility intervals associated with all variance components were wide for all random effects (Table 1).

**Fig 4.**
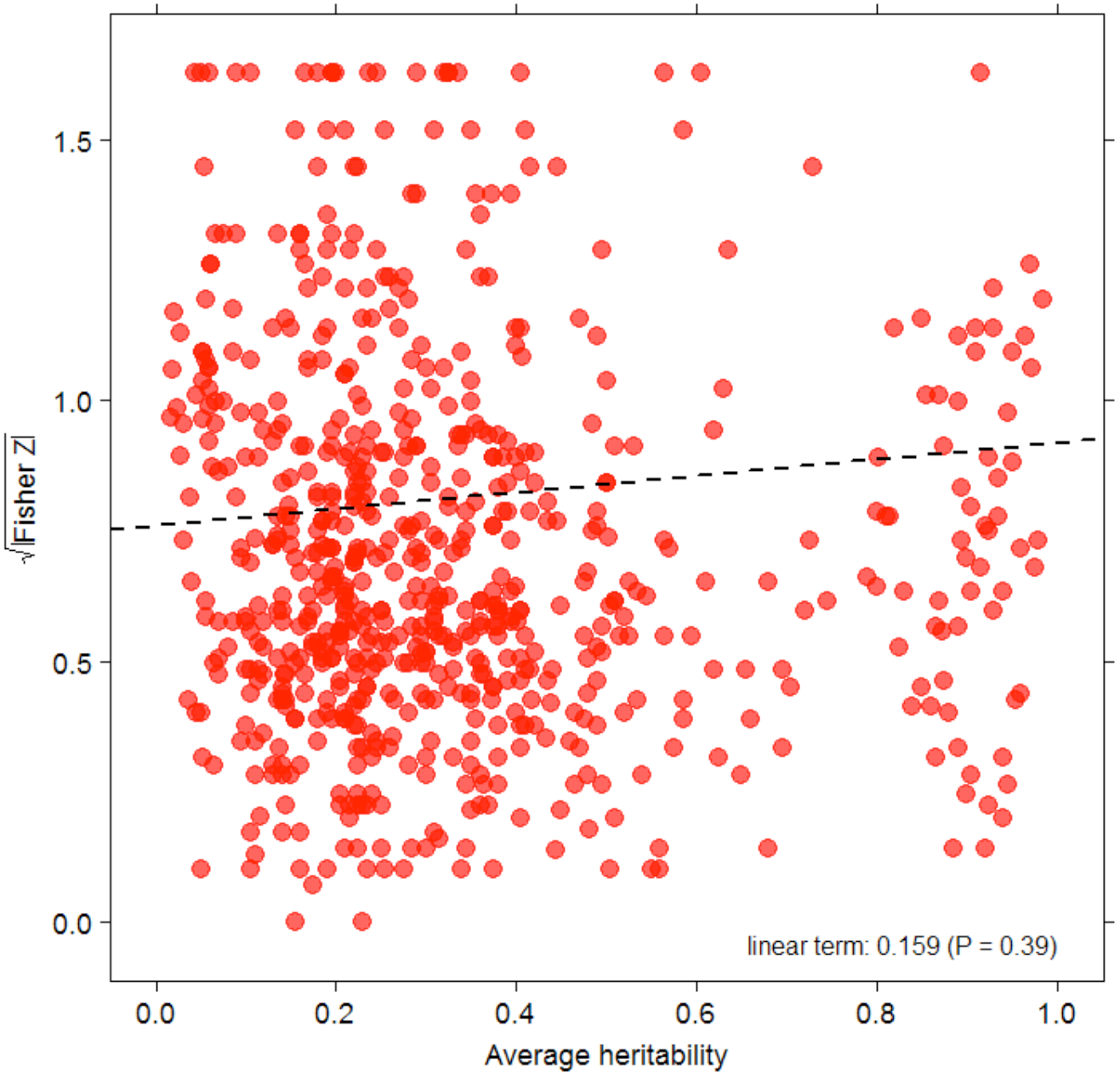
A relationship between absolute effect size (square-root of |Fisher Z|) and heritability averaged between-traits for cross-traits *r_g_*.

**Fig 5.**
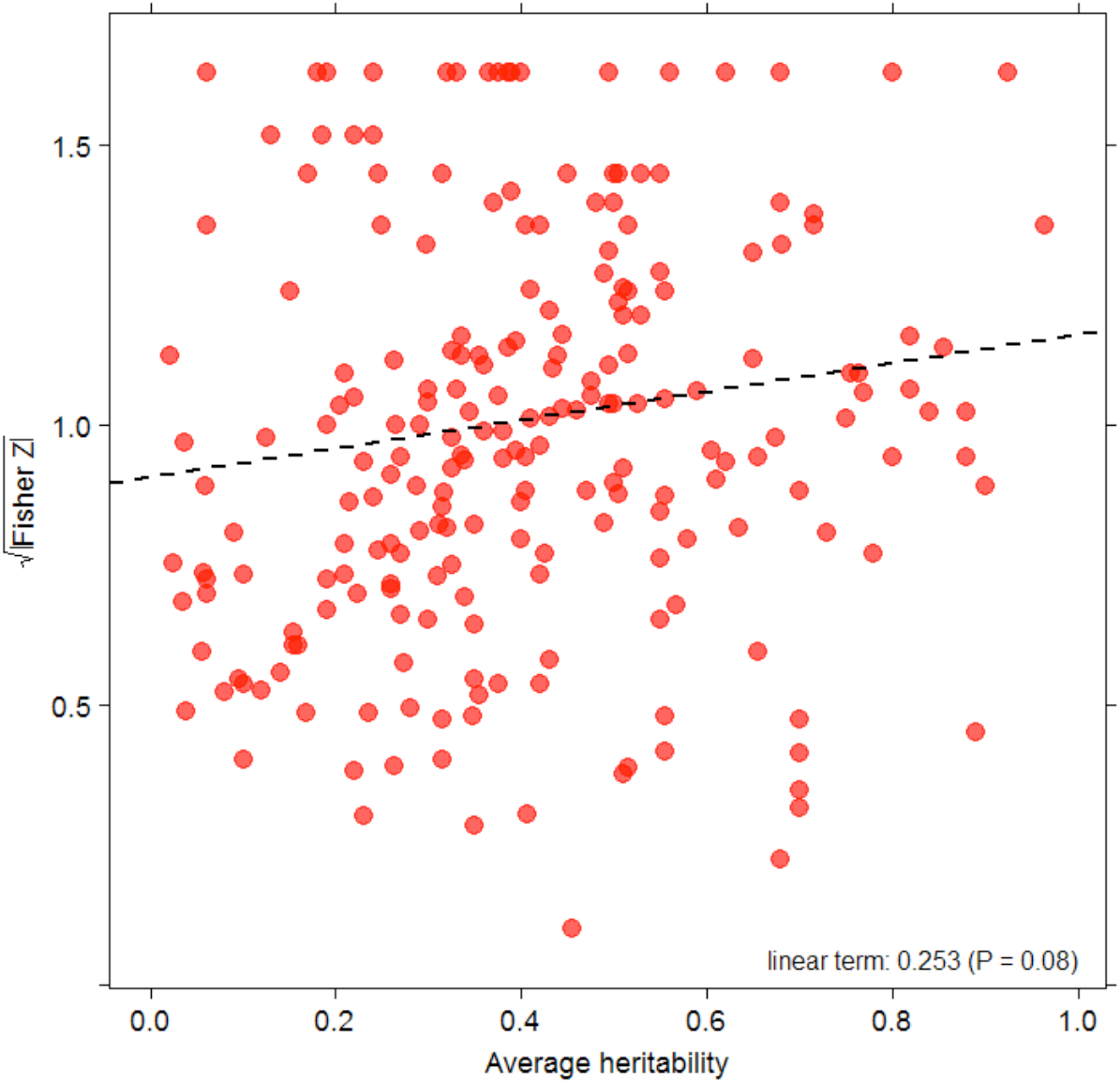
A relationship between absolute effect size (square-root of |Fisher Z|) and heritability averaged between-traits for cross-sex *r_g_*.

**Table 1.** Meta-analytical linear mixed models for the absolute effect size. Estimates are provided with their 95% credibility intervals. Upper parts of respective models contain fixed-effects estimates, lower parts provide variance components.

| Model term | Estimate | 95% CI | P |
| --- | --- | --- | --- |
| *Cross-sex $r_g$* |  |  |  |
| Intercept | 0.908 | (0.756; 1.062) | <0.0001 |
| Average $h^2$ | 0.253 | (-0.026; 0.518) | 0.08 |
| Difference of $h^2$'s | -0.055 | (-4.932; 0.405) | 0.79 |
| Taxon ID | 0.012 | (0; 0.037) | - |
| Study ID | 0.032 | (0.006; 0.064) | - |
| Residual | 0.034 | (0.021; 0.050) | - |
| *Between-trait $r_g$* |  |  |  |
| Intercept | 0.762 | (0.621; 0.907) | <0.0001 |
| Average $h^2$ | 0.159 | (-0.197; 0.521) | 0.39 |
| Difference of $h^2$'s | -0.465 | (-0.830; -0.092) | 0.02 |
| Taxon ID | 0.032 | (0; 0.072) | - |
| Study ID | 0.016 | (0; 0.053) | - |
| Trait 1 ID | 0.022 | (0.009; 0.036) | - |
| Trait 2 ID | 0.039 | (0.021; 0.056) | - |
| Residual | 0.015 | (0.0002; 0.037) | - |

Covariate expressing the difference between heritabilities appeared significant in case of between-trait *r_g_* meta-analysis: traits with larger differences between respective *h^2^* tended to have lower absolute Fisher’s *Z* (Table 1).

Analysis of the probability of obtaining *r_g_* significantly different from zero indicated, that it is positively associated with the average *h^2^* of underlying traits (GLMM, logistic regression coefficient 4.55, P=0.003; Fig. 6). However, even for the lowest average heritabilities predicted probability of obtaining significant *r_g_* estimate was 0.59 (Fig. 6).

**Fig. 6.**
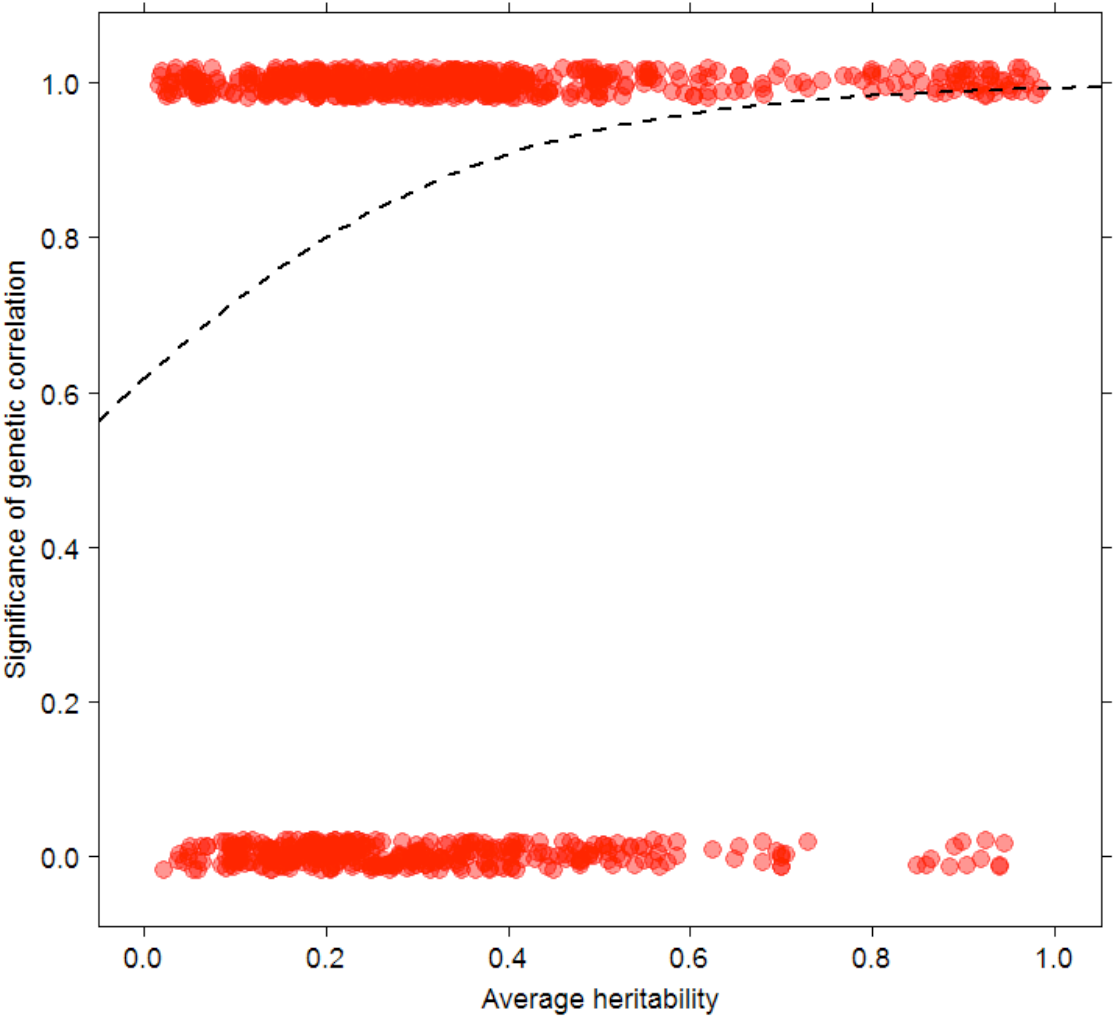
Relationship between obtaining significant estimate of *r_g_* (1 means significant) and average heritability. Dashed line depicts fitted logistic regression. Random noise along the y axis was introduced to data points to make reading of the plot easier.

Diagnostic funnel plots for both meta-analyses are presented in Fig. 7. In case of between-trait correlations, residual effect sizes form a symmetrical funnel indicating lack of publication bias (Egger’s regression slope: *b* = 0.003, *P* = 0.45). In case of cross-sex genetic correlations the funnel plot is less symmetrical, however Egger’s regression does not indicate any publication bias ( *b* = 0.02, *P* = 0.24). Inspection of residual variance from meta-analytical models indicates that inclusion of measurement error variance significantly lowered estimated residual variance, with the smallest change in residual variance observed in case of cross-sex *r_g_* meta-analysis (ordinary LMM: *V_R_* = 0.65 (0.53; 0.78); meta-analysis LMM: *V_R_* = 0.15 (0.02; 0.30)). It justifies the notion that measurement error variance in this case provides important extra information on the variance underlying examined relationship.

**Fig. 7.**
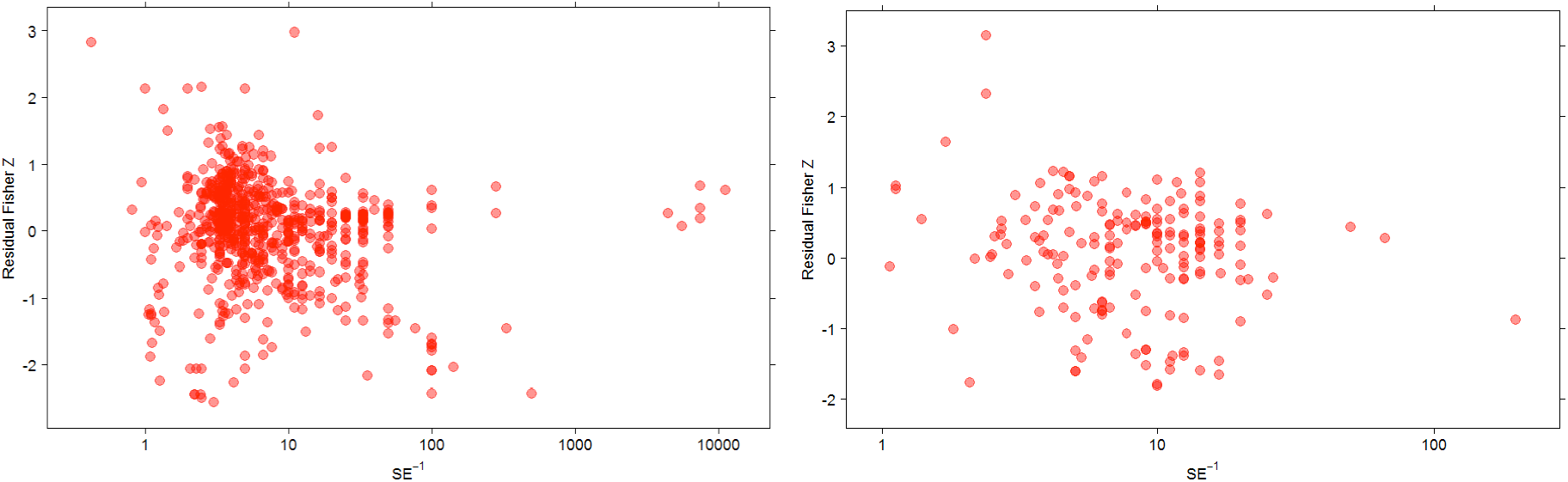
Funnel plots for between-traits (left) and cross-sex (right) genetic correlation analyses. Plots depict residual absolute effect sizes relative to the inverse of the standard error (depicted on logged scale for clearer presentation).

Simulated data indicate that in typical situations, i.e. when the separation of genetic effects is satisfactory and additive genetic variance actually estimates true genetic variance with little or no added variance, genetic correlations should not be associated with underlying heritabilities (Fig. 8, upper lines). However, strong positive relationship between *r_g_* and *h^2^* appears if additional variation is present in a substantial quantity compared to the magnitude of the true additive genetic variance. If such additional variation cannot be not separated from the genetic effect, strong bias arises in estimates of *rg*, generating strong *r_g_* – *h^2^* relationship. Depiction of estimated covariance ellipses (Fig. 9) clearly shows that with large amounts of non-genetic variance correlations between traits become blurred and more difficult to estimate.

**Fig. 10.**
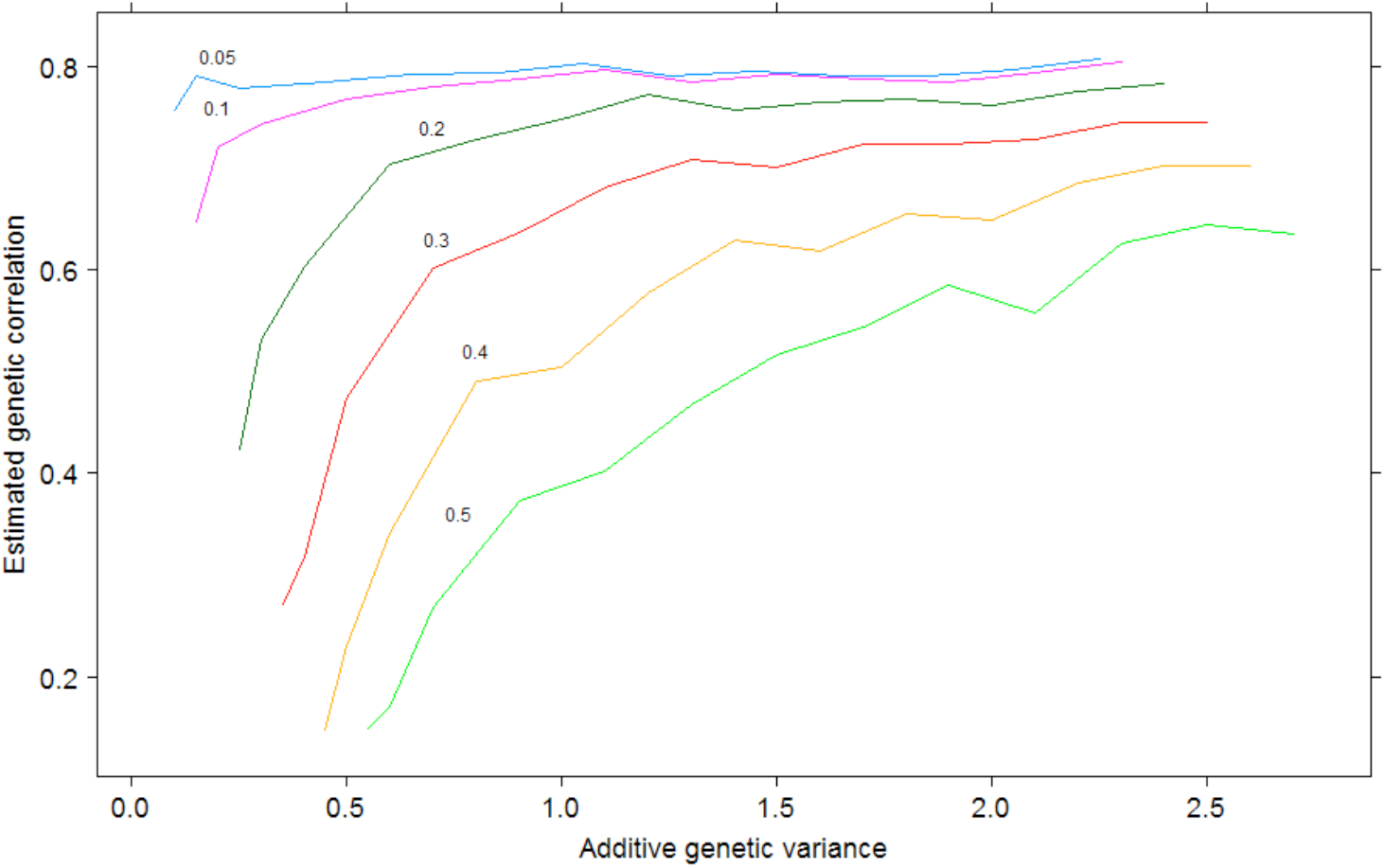
Results of simulation study of the dependence of estimated *r_g_* on the additive genetic variance values (V_A_ directly translates into heritabilities once residual variance is included). Numbers above lines provide amounts of additional, non-genetic variance used in each series of samplings.

**Fig. 11.**
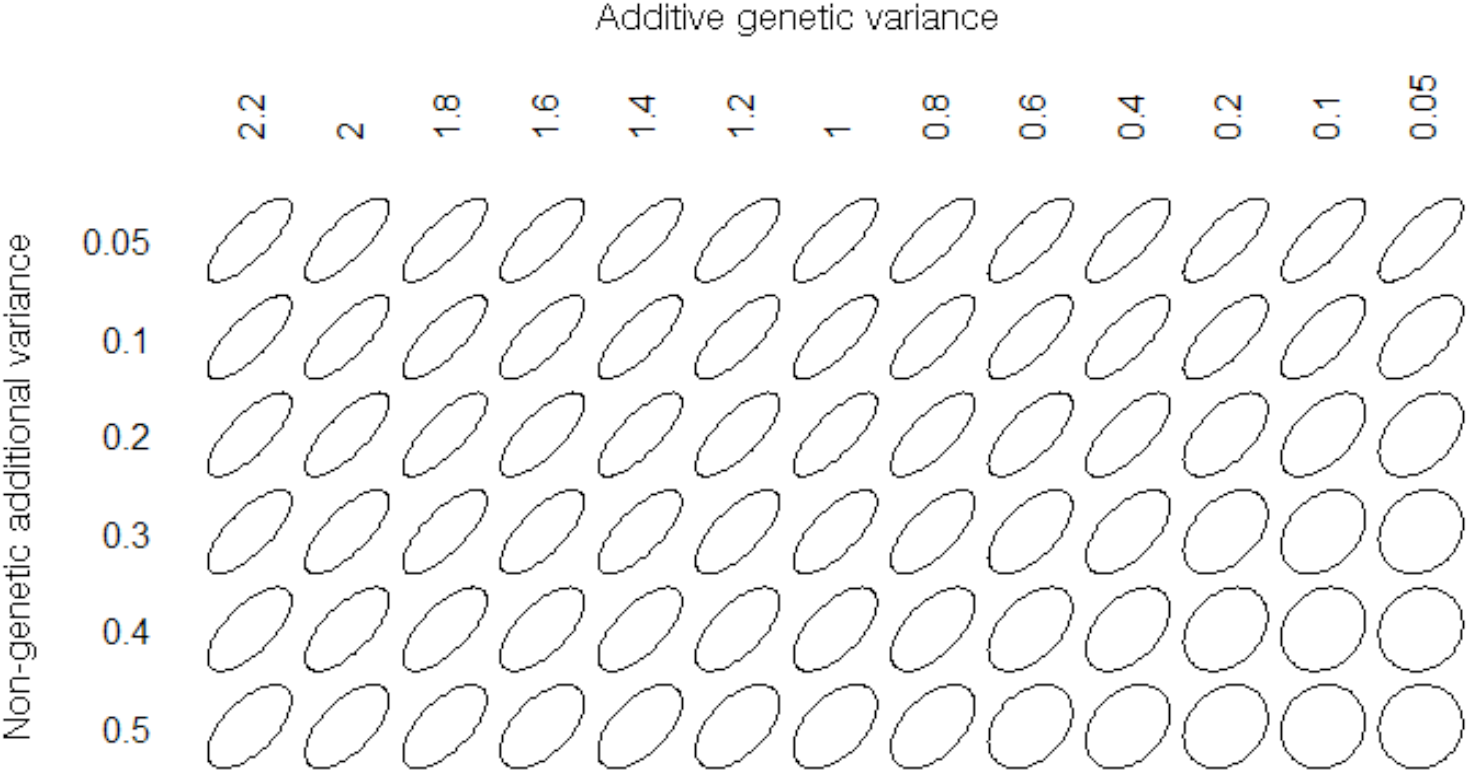
Covariance ellipses resulting from performed simulations, arranged on the grid of additive genetic variance values (horizontal axis) and non-genetic additional variance values (vertical axis) used to generate the data.

## Discussion

Here we show that the relationship between genetic correlations and heritabilities of underlying traits estimated using robust and powerful meta-analytical approach, employing large sample of published estimates, are weak and statistically non-significant, both for cross-traits and cross-sex genetic correlations. Approximately only 0.001% of the total variation in observed genetic correlations could be attributed to the direct relationship with heritability. Thus, our findings challenge widely accepted belief that low heritabilities are associated with low and non-significant or less detectable genetic correlations.

The theory of quantitative genetics offers several possible mechanisms that could generate such relationship. The most important candidate explanations include (i) higher environmental variability in life-history and fitness traits (associated with lower heritabilities) (Price and Schluter 1991; Merilä and Sheldon 1999), (ii) biased estimates of *r_g_* due to selection acting on one of the traits (Brown 1969; Lynch and Walsh 1998) and (iii) statistical issues leading to the incomplete separation of genetic effects from other sources of phenotypic variation (most importantly, common environment and parental effects). Our meta-analysis does not support the first two explanations. In both analyzed datasets the type of the trait had no influence on the strength and direction of *r_g_* – *h^2^* relationship, as indicated by non-significant trait type vs. linear slope interactions. Similarly, lack of any significant interactions between the linear slope and type of the study (agricultural, laboratory and field) provides an indirect evidence against the selection-related bias: one could expect that agricultural studies focus on individuals strongly selected with respect to valuable productivity traits and hence lack of heterogeneity in slopes across study types excludes selective explanation. It is more difficult to relate our results to statistical problems with decompositions of different sources of phenotypic variance. Linear models theory developed a range of methods capable of separating different sources of variation in phenotypic traits (Kruuk and Hadfield 2007). Precision and robustness of these methods differs considerably, especially with respect to genetic correlations (as they require larger sample sizes): most accurate and robust are methods using available pedigree information (“animal model”) (Kruuk 2004; Kruuk and Hadfield 2007), followed by half-sib and full-sib approaches. However, lack of significant interactions between slopes of analyzed relationships and statistical methodology applied does not provide support for this explanation. Simulation study performed along with this meta-analysis confirmed that in the presence of large additional variance, not separated from additive genetic effects, decreasing genetic variance (and hence – heritability) may lead to a downward bias in the estimates of *r_g_*. In this context it appears that the degree to which published data violate the assumption of complete separation of genetic effects seems to be negligible.

Statistical sampling issues may have profound effects not only on the value but also detectability of *r_g_*. By detectability we understand the likelihood that under certain heritability values the method employed is able to yield statistical significance at the assumed confidence level (here we assumed confidence equal to 95%). As shown above, genetic correlations are not significantly related to heritabilities and hence detectability of *r_g_* should not be lower for low values of *h^2^*. Surprisingly, we have shown that detectability of *r_g_* is weakly positively related to heritability. However, even for the lowest heritabilities, the predicted probability of obtaining significant estimates of *r_g_* is not lower than 50%. Hence, we conclude that detectability is not substantially compromised by low values of heritabilities.

Apart from sampling issues, our results have other profound statistical consequences. Genetic correlations in our meta-analysis were subdivided onto two distinct groups: between-traits and cross-sex genetic correlations. Cross-trait genetic correlations are usually studied when looking for genetic trade-offs and constraints between phenotypic traits (e.g. Norry et al. 2000). Such studies usually assume that the null hypothesis is in the form H_0_: *ρ_g_* = 0, which is reasonable taken that under random mating two traits should not share any genetic background (excluding cases of pleiotropy) and hence, should not be genetically correlated (Lynch and Walsh 1998). In other words, any studies analyzing low-heritability traits would only risk elevated levels of type II error (false negatives) if the positive *r_g_* – *h^2^* correlation was actually true. However, testing the cross-sex genetic correlations usually assumes the null hypothesis of the form H_0_: *ρ_g_* = 1, which is logical as we expect full genetic correlation of individuals carrying identical sets of genes and differing only with their sex (Robertson 1959; Eisen and Legates 1966; once again we exclude special cases of sex-linked traits). In this case, analyses of low-heritability traits would be associated with elevated type I error rates (falsely significant results) in the presence of the significant positive *r_g_* – *h^2^* correlation. In this context our results do not justify the fact that cross-sex genetic correlations of weakly heritable traits are associated with higher levels of statistical error.

An important feature of the presented relationships between *r_g_* and average *h^2^* is that it might be influenced by publication bias present in the data. Such bias could be expected as strong genetic correlations are more attractive in case of cross-trait studies, the opposite being true in the case of cross-sex r_g_’s where low correlations are considered more interesting. However, diagnostic plots indicate no publication bias in both analyzed datasets.

The only significant correlation found in our meta-analysis was detected for the relationship between *r_g_* and the difference between heritabilities in case of between-traits genetic correlations. It appears that lower genetic correlations are associated with more extreme values of heritabilities. Explaining this pattern is difficult. As over 95% of all data-points had these differences lower than 0.3, it is possible that this result is a statistical artifact arising due to strongly asymmetrical distribution of differences. Clearly, further exploration of this pattern is required.

To conclude, our review and meta-analysis of published estimates of genetic correlations and heritabilities does not support the prediction that genetic correlations are strongly positively associated with heritabilities of underlying traits. Thus, we think that the criterion of low heritability should not be treated lightly and too confidently when evaluating the quality and reliability of published estimates of genetic correlations.

## Acknowledgements

This study was supported by the grant of the National Science Center number NN304061140 (to Sz.D.) and by the START program of the Foundation for Polish Science (to Sz.D.)

